# Lost in translation: the pitfalls of Ensembl Gene annotations between human genome assemblies and their impact on diagnostics

**DOI:** 10.1101/2020.11.12.380295

**Authors:** Mohammed O.E Abdallah, Mahmoud Koko, Raj Ramesar

## Abstract

**Background:** The GRCh37 human genome assembly is still widely used in genomics despite the fact an updated human genome assembly (GRCh38) has been available for many years. A particular issue with relevant ramifications for clinical genetics currently is the case of the GRCh37 Ensembl gene annotations which has been archived, and thus not updated, since 2013. These Ensembl GRCh37 gene annotations are just as ubiquitous as the former assembly and are the default gene models used and preferred by the majority of genomic projects internationally. In this study, we highlight the issue of genes with discrepant annotations, that have been recognized as protein coding in the new but not the old assembly. These genes are ignored by all genomic resources that still rely on the archived and outdated gene annotations. Moreover, the majority if not all of these discrepant genes (DGs) are automatically discarded and ignored by all variant prioritization tools that rely on the GRCh37 Ensembl gene annotations.

**Methods:** We performed bioinformatics analysis identifying Ensembl genes with discrepant annotations between the two most recent human genome assemblies, hg37, hg38, respectively. Clinical and phenotype gene curations have been obtained and compared for this gene set. Furthermore, matching RefSeq transcripts have also been collated and analyzed.

**Results:** We found hundreds of genes (N=267) that were reclassified as “protein-coding” in the new hg38 assembly. Notably, 169 of these genes also had a discrepant HGNC gene symbol between the two assemblies. Most genes had RefSeq matches (N=199/267) including all the genes with defined phenotypes in Ensembl genes GRCh38 assembly (N=10). However, many protein-coding genes remain missing from the current known RefSeq gene models (N=68)

**Conclusion:** We found many clinically relevant genes in this group of neglected genes and we anticipate that many more will be found relevant in the future. For these genes, the inaccurate label of “non-protein-coding” hinders the possibility of identifying any causal sequence variants that overlap them. In addition, Important additional annotations such as evolutionary constraint metrics are also not calculated for these genes for the same reason, further relegating them into oblivion.

## Background

### Ensembl Gene Annotation Gap

The human reference genome has been continually updated since the release of the first draft human genome. The current release, GRCh38, released in 2013, is the most complete and only gap-free human genome assembly so far (1). Despite all its imperfections, the previous human genome assembly GRCh37 (also known as hg19) which was released in 2009 is still the predominant human genome assembly in current use. An obvious explanation for this lag in adopting the newest assembly is the availability of vast amounts of relational genomic annotations for the previous assembly that are not yet ported to the current one, as well as the incurred cost of remapping or recalling genomics data under the new assembly. We previously raised the issue of minor reference alleles in the GRCH37 assembly, how it adversely impacts variant calling, and its implications especially for recessive genetic disorders (2). In this correspondence we highlight another pressing issue that is related to GRCh37 annotations.

A critical aspect of genomic analysis whether in the research or diagnostic setting is the choice of gene models. Gene models delineate gene coordinates in the assembly as well as various exon-intron and coding sequence (CDS) boundaries. More importantly, it also determines gene biotypes: assigning which genes are protein-coding vs pseudogenes, etc. Different gene models exist in genomics research today, however, most studies either use the NCBI RefSeq (3,4) or EBI Ensembl gene-models. The Ensembl gene models - based on Gencode (5)-are the most comprehensive and well-curated gene models currently available and boast more genomic coverage than its RefSeq counterpart (6). No wonder then it has become the default gene model for almost all the major genomics projects including Encode (7,8),the 1000 Genomes project (9,10), the 100K genomes project (Genomics England) (11), gnomAD (12) and many others.

A quick survey of the major projects listed above reveals that all of them rely on the old GRCh37 (based on the Gencode v19) version of Ensembl genes. However, as we discuss in this letter, there is a major issue with this situation, namely, the lack of any updates to this version, which has been archived since 2013. Thus, hundreds of protein-coding genes that have been added or updated since then have not been backported to this widely used version of Ensembl genes. We will hereafter refer to these genes as discrepant genes (DGs). There are many genes that are classified as protein-coding in the new version but used to be classified as non-protein-coding in the old version. This issue is further compounded by the fact that many of these genes also have new HGNC (13) names and are not easily traceable to their previous HGNC gene names, effectively creating an annotation limbo for potentially significant genes.

## Results

To identify genes with discrepant annotations, we assessed the biotype annotations for Ensembl genes between the two assemblies, we found hundreds of genes (N=267) that were reclassified as “protein-coding” in the new hg38 assembly. Notably, 169 of these genes also had a different HGNC gene name between the two assemblies, further complicating the mission of how to trace them effectively between assemblies: searching the Ensembl GRCh37 website or any of the other resources that depend on the GRCh37 genes with the new HGNC names does not yield any results linking the two gene name.

Interestingly, we also found that 10 of those DG have known clinical phenotypes. Six of those have shared phenotypes between the two assemblies while four of them have new phenotype annotations exclusively in the new assembly. Still, those new phenotypes could not be found in the GRCh37 assembly. A table of the phenotypes and their discrepancies between the two assemblies is summarized in Table 1.

**Table 1.**
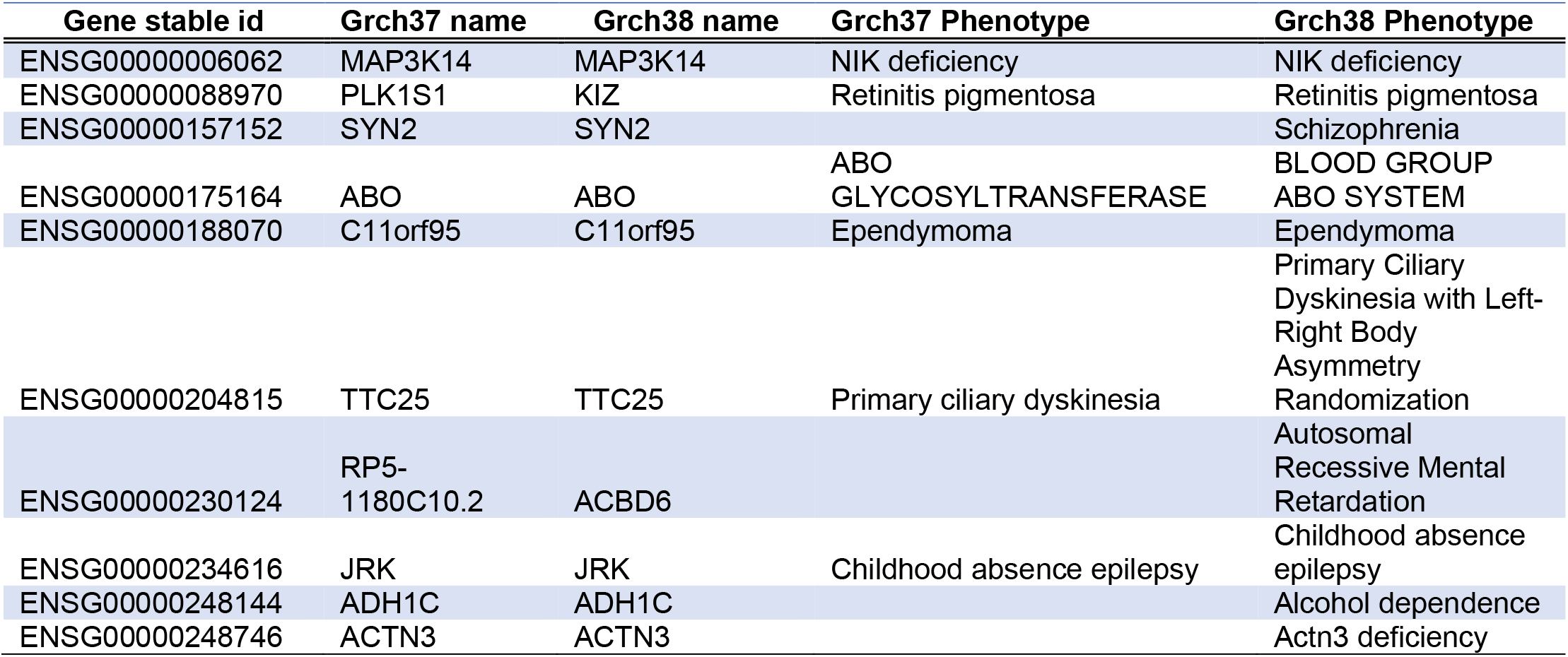

We also evaluated if using RefSeq gene models instead of the Ensembl genes would solve or alleviate this annotation mismatch issue. Most genes had RefSeq counterparts (N=199/267) including all the genes with defined phenotypes in Ensembl genes GRCh38 assembly (N=10). However, many protein-coding genes remain missing from the current known RefSeq gene models (N=68). An illustration of how DGs map to different categories (particularly, how many DGs have RefSeq transcripts or have a phenotype) is depicted in Figure 1.

**Fig 1.**
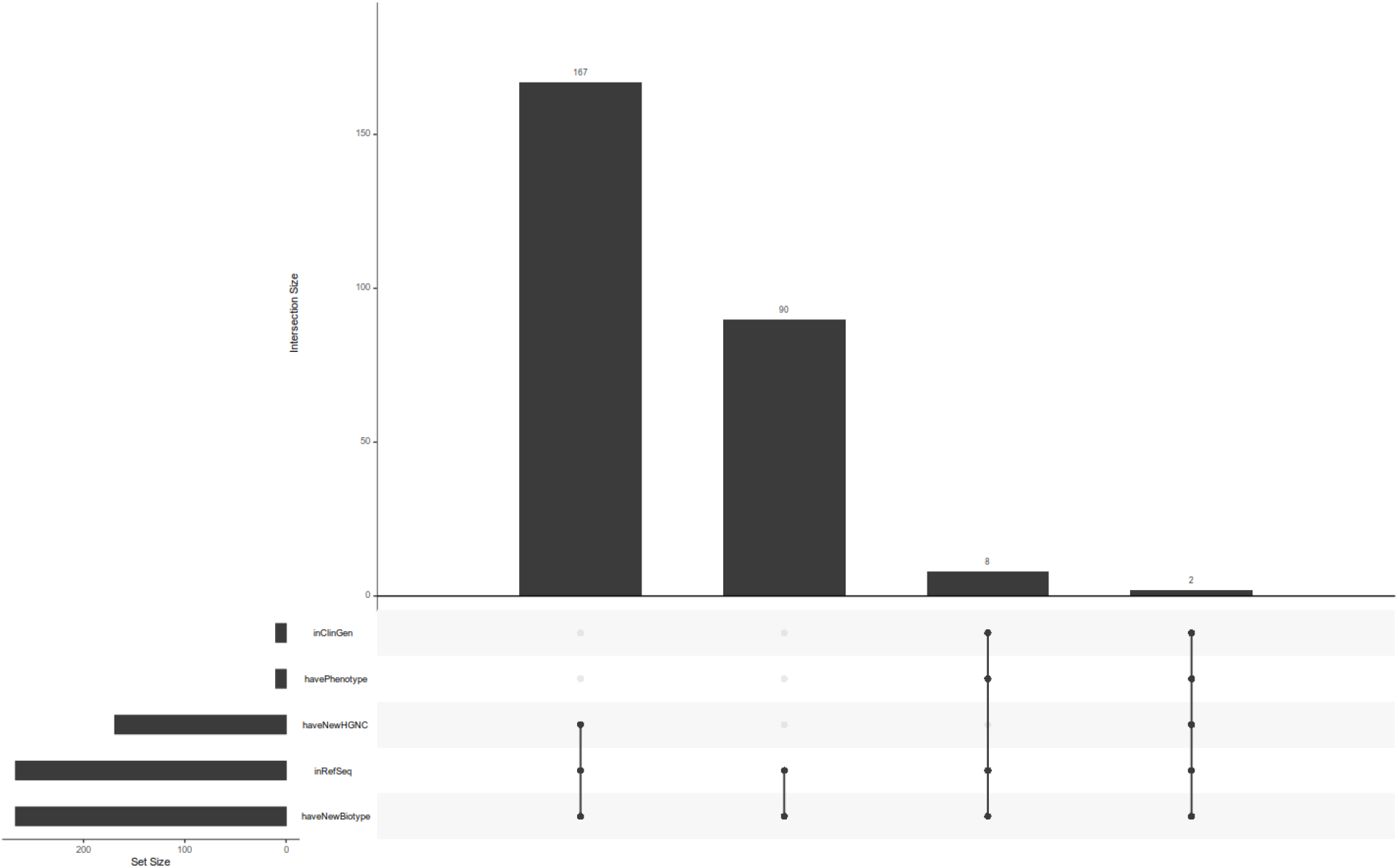
an UpSet diagram of DGs (discrepant genes) and their intersection with different annotation categories.

## Discussion

Despite their prevalent use in genomics today, the archived non-updated set of GRCh37 Ensembl genes has many overlooked pitfalls. The most glaring issue is the lag in assigning the correct biotype to hundreds of potentially causal genes.

Today, a sizable proportion of Mendelian genetic disorders eludes a definitive molecular diagnosis; having accurate and up to date gene models is a critical prerequisite to minimize the potential of genetic disorders remaining unlinked to their genes. Unfortunately, this pitfall is expected to remain unbridled as the use of GRCh37 data is prevalent and many of the major genomic analysis tools rely on this resource for downstream analysis. These commonly used genomic analytic tools including Exomizer, Gemini, OpenCGA, etc, automatically filter out ‘non-protein-coding genes’, thereby discarding potentially causal genes.

In addition to the aforementioned variant prioritization problem, gene prioritization and curation efforts for diagnostic gene panels is similarly affected. An example of this issue is the KIZ gene which is implicated in some forms of retinitis pigmentosa (previously known as PLK1S1 in the GRCh37 Ensembl gene model). Gene curation for this gene’s involvement in retinal disorders demonstrates such confusion. Interrogating the curation activity for this gene in the Genomics England PanelApp shows that it was excluded from the diagnostic retinal disorders gene panel because of the Ensembl GRCh37 misclassified biotype of “non-coding” gene. It was only after we engaged with the PanelApp curators and pointed out the unfortunate annotation issue with the KIZ gene that it was reassessed and incorporated into the diagnostic gene panel for retinal disorders (14).

Another issue that stems from this annotation gap is the lack of evolutionary constraint measures for these genes. The evolutionary constraint metrics have recently emerged as an important tool in our genomics arsenal. Loss-of-function and missense constraint are widely used to identify genes with a potentially mendelian signature of constraint. However, querying the ExAC or gnomAD resources for any of the DGs will return no results for constraint as both resources erroneously discard those genes as non-protein-coding. Even more recent attempts to map the constrained coding regions of the human genome failed to include these DGs regions as it was again filtered out as non-coding genes (15).

We also found that although using RefSeq genes instead of the Ensembl GRCh37 genes would solve many variant prioritization issues for hg19 genomic data, not all protein coding genes are present in RefSeq known genes (RefSeq NM). Furthermore, There is still a wide chasm between the two gene models, although the new Matched Annotation from NCBI and EMBL-EBI (MANE) project (16) will help this bridge the gap in the future.

## Conclusion

Our results illustrate the need for up to date annotations and careful examination of gene curations for genomic or clinical diagnosis of genetic disorders. We highlighted many ensuing issues that stem from the annotations gap in the widely used Ensembl gene models including issues with variant and gene prioritization. We recommend testing genomic data against the new GRCh38 assembly and/or the RefSeq gene models for better diagnostic performance and accuracy.

## Methods

### Finding genes with discrepant annotations and their phenotypes

We utilized Ensembl Biomart to find all gene and gene annotations from both the GRCh37 and GRCh38 assemblies and extracted all discrepant genes (DGs) that have the updated biotype of protein-coding in the new assembly but not the old assembly. We further investigated their associated phenotype from multiple resources including Biomart (17,18), Clinvar (19), GWAS Catalog (20), and CellBase (21).

### Gene Disease Curation

To determine the clinical utility of DGs and investigate their clinical curation status, we traced their current gene-disease curation using the industry-standard ClinGen gene curation interface and the more active Genomics England PanelApp.The latter is a crowdsourced gene curation resource that branched out of the 100K genome project and has considerably more gene curations than the ClinGen resource (22) and is also part of the Gene Curation Coalition. Both resources pool information from other approved resources including OMIM (23), and Orphanet (24).

### RefSeq Mapping

We also investigated All discrepant Ensembl genes against their RefSeq counterparts using the grpofiler2 R package (25), specifically, we used the gConvert component of gProfiler (26) to perform the id mapping between Ensembl genes and their RefSeq counterparts. We also noted that using Ensembl Biomart service yielded incomplete mapping between Ensembl and RefSeq genes.

## Declarations

### Ethics approval and consent to participate

Not applicable.

### Consent for publication

Not applicable.

### Availability of data and materials

The datasets analyzed in this study are available publicly from Ensembl Biomart and gProfiler. The R script used in the analysis is publicly available in github and is completely reproducible from https://github.com/melsiddieg/EnsGenes.

### Competing interests

The authors declare no competing interests.

### Funding

Not applicable.

### Authors’ contributions

MOEA and RR conceived the idea. All authors contributed to the design. MOEA and MK performed the bioinformatic analysis. All authors contributed to the interpretation of findings. The manuscript was written by MOEA with intellectual input and critical revisions from MK and RR All authors read and approved the final manuscript.

## Acknowledgements

Not applicable.

## Abbreviations

## Notes

https://github.com/melsiddieg/EnsGenes

### Competing Interest Statement

The authors have declared no competing interest.

